# Transcriptional remodeling of ubiquitin regulatory networks during trained immunity

**DOI:** 10.64898/2026.05.06.723281

**Authors:** John Santelices, Zoe Schaefer, Wambui Gachunga, Cameron Celeste, Ivana K. Parker

## Abstract

**Background:** Trained immunity is a durable functional reprogramming of innate immune cells characterized by enhanced responsiveness upon secondary challenge. While metabolic rewiring and epigenetic remodeling are well-established features of this process, the contribution of ubiquitin-mediated post-translational regulation remains poorly defined.

**Methods:** We performed an integrative analysis of publicly available human transcriptomic datasets derived from monocytes, macrophages, and PBMCs exposed to established training stimuli (β-glucan, Bacillus Calmette–Guérin [BCG], and hemin–β-glucan) followed by secondary stimulation. A curated panel of deubiquitinating enzymes (DUBs) and E3 ubiquitin ligases with established immune functions was analyzed for differential expression. Gene Ontology (GO) and KEGG pathway enrichment analyses were conducted to evaluate higher-order convergence across independent datasets.

**Results:** Across multiple trained immunity models, we identified reproducible transcriptional remodeling of ubiquitin-modifying enzymes. USP25, OTUB1, and TRIM25 were consistently upregulated following restimulation, whereas several chromatin- and cytokine-regulatory DUBs—including USP3, USP4, USP7, USP16, MYSM1, and USP38—were downregulated. Normalization to RPMI-restimulated controls reduced many activation-associated signals; however, USP25 remained persistently elevated, suggesting a stable training-associated signature. Pathway enrichment analysis independently demonstrated significant engagement of ubiquitin-related functional categories across datasets, supporting coordinated reorganization of ubiquitin regulatory networks.

**Conclusion:** These findings identify selective transcriptional remodeling of the ubiquitin– proteasome system as a recurring feature of trained immunity. Integrating ubiquitin signaling into the established metabolic–epigenetic framework expands the conceptual model of innate immune memory and suggests that ubiquitin-modifying enzymes function as modulatory rheostats shaping immune amplitude and stability. Future functional and proteomic studies are required to determine whether these transcriptional signatures directly mediate trained immunity phenotypes.

## Introduction

Trained immunity describes the long-term functional reprogramming of innate immune cells following exposure to specific stimuli, resulting in an altered — often enhanced — response to subsequent stimulation after cells return to a resting state [1], [2]. Unlike immunological tolerance, which dampens responsiveness, trained immunity increases secondary responses and represents a durable form of innate immune adaptation driven by epigenetic reprogramming rather than gene recombination [1]. Well-characterized inducers include β-glucan (βG), lipopolysaccharide (LPS), and the bacillus Calmette–Guérin (BCG) vaccine [2].

The persistence of trained immunity is governed by epigenetic remodeling, particularly histone modifications associated with active chromatin. Key marks include histone 3 lysine 27 acetylation (H3K27ac) at distal enhancers, often coupled with histone 3 lysine 4 methylation (H3K4me1), and histone 3 lysine 4 trimethylation (H3K4me3) at gene promoters [2]. These chromatin changes are tightly linked to metabolic rewiring, as cellular metabolism regulates the availability of metabolites required by chromatin-modifying enzymes, coupling immunometabolism to innate immune memory [3], [4].

The ubiquitin–proteasome system (UPS) also plays a vital role in immune signal transduction [5], [6]. E3 ubiquitin ligases and deubiquitinating enzymes (DUBs) dynamically control protein stability and function through reversible ubiquitin modifications, influencing processes ranging from proteasomal degradation to signaling complex assembly [7]. These modifications regulate pathways critical for innate immune activation and immunometabolism, including glycolysis and NF-κB signaling [7] - [10]. Beyond signaling, ubiquitination also shapes chromatin architecture: polyubiquitination can promote histone degradation, while monoubiquitination coordinates crosstalk with histone methylation and acetylation [11]. For example, phosphorylation at histone 3 threonine 11 (H3T11) triggers EGFR-mediated K48-linked polyubiquitination at H3K4, leading to histone turnover [12], whereas monoubiquitinated H3 recruits DNMT1 to DNA, promoting cytosine methylation [13], [14]. Both DNMT1 and histone ubiquitination are regulated by the DUB USP7, a clinically investigated cancer therapeutic target [15] - [17].

Although ubiquitin signaling is essential to immune regulation, its specific contribution to trained immunity remains largely undefined. In this study, we analyze multiple publicly available *in vivo* and *ex vivo* trained immunity gene expression datasets to evaluate whether known ubiquitin-modifying enzymes, including DUBs and E3 ligases, are transcriptionally altered following trained immunity induction and secondary stimulation. Our integrative analysis revealed a distinct expression signature characterized by consistent upregulation of USP25, OTUB1, and TRIM25, alongside suppression of several chromatin- and cytokine-regulatory DUBs across multiple trained immunity models. These findings suggest that ubiquitin signaling may represent an underappreciated regulatory layer of innate immune memory and provide a foundation for future mechanistic studies into how ubiquitin-dependent post-translational control intersects with metabolic and epigenetic reprogramming in trained immunity.

## Results

### Ubiquitin-modifying enzymes are differentially expressed across trained immunity datasets

To investigate whether the ubiquitin–proteasome system contributes to trained immunity regulation, we analyzed publicly available transcriptomic datasets derived from monocytes, macrophages, and peripheral blood mononuclear cells (PBMCs) exposed to established training stimuli, including β-glucan, Bacillus Calmette–Guérin (BCG), and hemin–β- glucan, followed by secondary stimulation (**Table 1**). Expression profiles of deubiquitinating enzymes (DUBs) and E3 ubiquitin ligases with known roles in immune signaling were examined alongside canonical trained immunity markers and cytokines for context (**Supplementary Tables 1–3**).

**Table 1.**
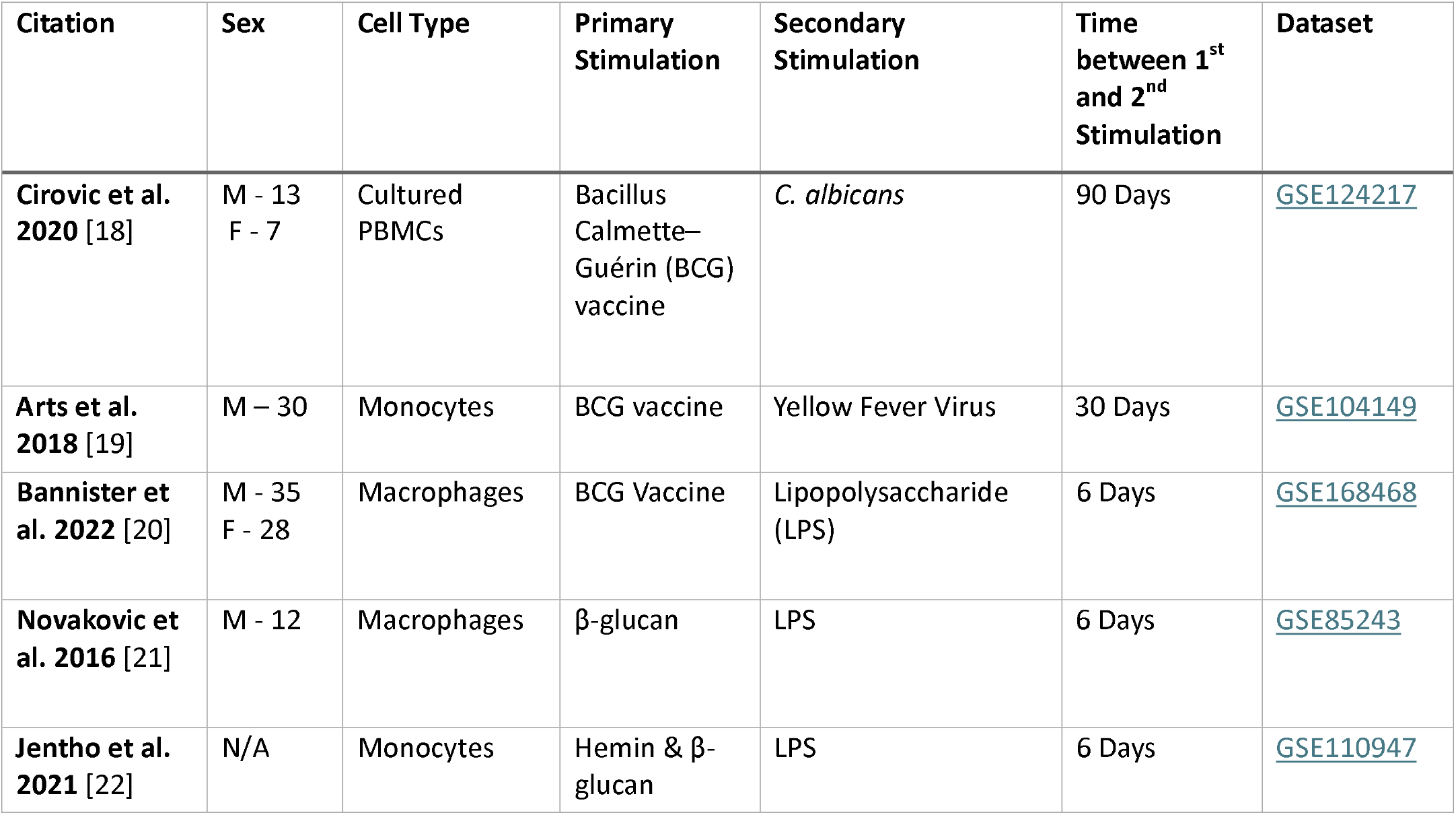
A description of gene data sets used for this analysis with associated PubMed ID number, experimental cell type, primary stimuli, secondary stimuli, time between stimulations, citation, and GEO data links.

Across independent datasets, a reproducible pattern of transcriptional remodeling was observed within the ubiquitin machinery. The DUBs USP25 and OTUB1, together with the E3 ligase TRIM25, were consistently upregulated following restimulation relative to baseline controls (**Figure 1**). These enzymes are established regulators of antiviral signaling and inflammatory balance, acting on key intermediates such as TRAF3, TRAF6, and RIG-I. Notably, USP25 exhibited stable elevation across cell types, stimuli, and time points, suggesting a broadly conserved transcriptional response [18] - [22].

**Figure 1.**
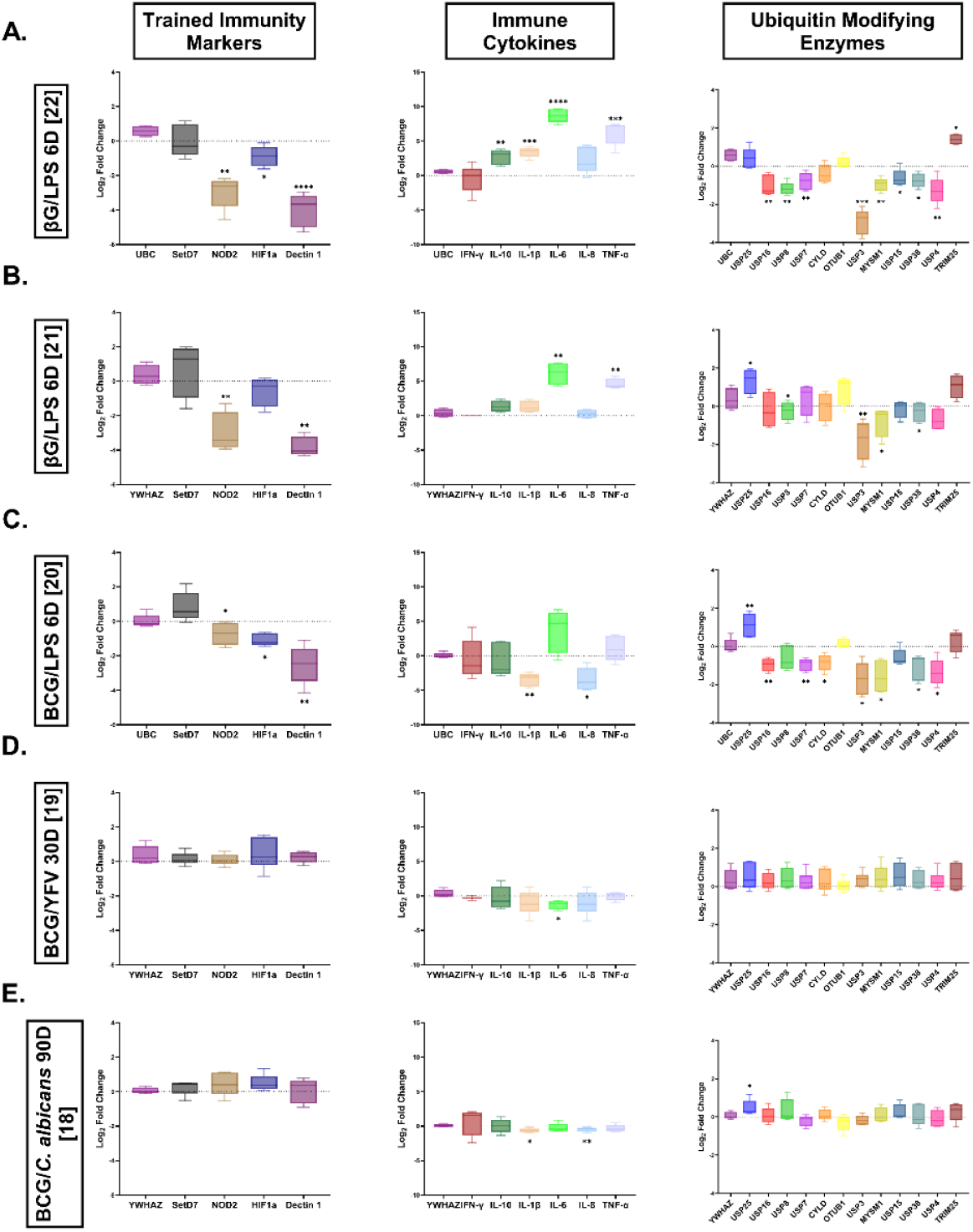
Gene-level remodeling of trained immunity markers, immune cytokines, and ubiquitin-modifying enzymes across independent trained immunity datasets. Panels (A–E) represent distinct trained immunity models as indicated along the left margin: (A) β-glucan (βG) monocytes, 6-day rest, LPS restimulation [22]; (B) β-glucan macrophages, 6-day rest, LPS restimulation [21]; (C) BCG macrophages, 6-day rest, LPS restimulation [20]; (D) BCG monocytes, 30-day rest, YFV restimulation [19]; (E) BCG PBMCs, 90-day rest, *C. albicans* restimulation [18]. Within each panel, box plots display log2 fold change in gene expression relative to baseline controls for trained immunity markers (left column), immune cytokines (middle column), and ubiquitin-modifying enzymes (right column). Each box represents donor replicates within the respective dataset. Statistical significance was determined using parametric paired t-tests (p < 0.05). Collectively, panels illustrate reproducible yet selective transcriptional remodeling across independent training conditions.

USP25 has previously been reported to protect TRAF3 and TRAF6 from proteasomal degradation, thereby dampening excessive pro-inflammatory cytokine release while promoting type I interferon production [23] - [25]. Likewise, TRIM25 and OTUB1 are established regulators of RIG-I activation, coordinating ubiquitination and deubiquitination to enhance antiviral signaling [26] - [30].

In contrast, several chromatin- and cytokine-regulatory DUBs—including USP3, USP4, USP7, USP16, MYSM1, and USP38—were reproducibly downregulated after secondary challenge (**Figure 1A-1C & Figure 2**). These enzymes normally constrain NF-κB signaling or regulate histone ubiquitination states, indicating that their suppression may transiently relieve transcriptional restraint during trained immune activation (**Table 2**) [31] - [42].

**Table 2.** Summary of ubiquitin-modifying enzymes differentially regulated across trained immunity datasets. Table highlights selected ubiquitin-modifying enzymes identified as significant and reproducibly regulated in Figure 2. Arrows indicate the predominant direction of regulation across datasets (green = upregulated; red = downregulated). Primary molecular targets and established immune functions are included to contextualize potential roles in immune signaling and NF-κβ regulation. Listed genes represent consistent patterns observed across independent trained immunity models.

| Protein Name |  | DUB or E3 Ligase | Primary Targets | Immune Role |
| --- | --- | --- | --- | --- |
| USP25 | ↑ | DUB | TRAF3/6 | Type 1 interferon (IFN) response. Reduce proinflammatory cytokine production. |
| TRIM25 | ↑ | E3 Ligase | TRAF2/6<br>RIG-I | Ubiquitinates RIG-I to enhance RNA virus signaling and modulate TRAF2/6 signaling. |
| OTUB1 | ↑ | DUB | UBC13<br>RIG-I | Promotes NF-κB signaling by stabilizing UBC13. Activates RIG-I-dependent antiviral signaling. |
| USP3 | ↓ | DUB | RIG-I<br>MyD88 | Negative regulator for type I IFN signaling and NF-κB activation. Attenuates proinflammatory and antiviral response. |
| USP4 | ↓ | DUB | RIG-I<br>TRAF6 | Positive regulator of RIG-I activity and enhances NF-κB activation by targeting TRAF6. |
| USP7 | ↓ | DUB | MyD88<br>Raf-1 | Stabilizes a critical adaptor protein for NF-κB activation. Regulates ERK1/2 signaling pathway. |
| USP16 | ↓ | DUB | IKKβ<br>Histone H2A | NF-κB pathway induction and T-cell activation. |
| USP38 | ↓ | DUB | TBK1<br>KDM5B | Regulates H3K4 trimethylation and proinflammatory activity. |
| MYSM1 | ↓ | DUB | STING<br>RIP2<br>TRAF3/6 | Regulates antiviral and proinflammatory immune response through interaction with TRAFs, STING, and RIP2. |

**Figure 2.**
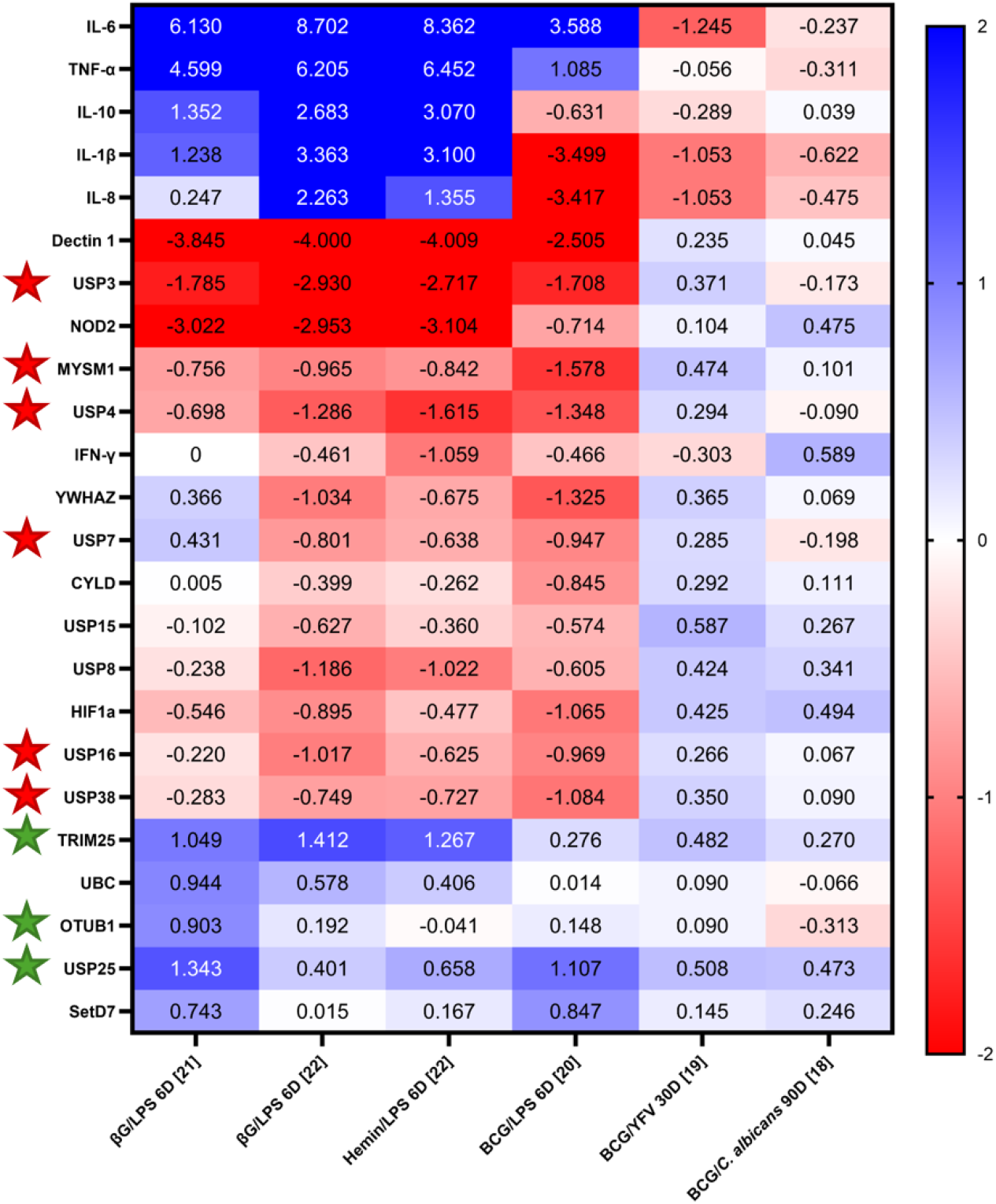
Heatmap of average log2 fold change in gene expression across trained immunity datasets relative to baseline controls. Rows represent individual genes and columns represent independent trained immunity datasets with indicated training and restimulation conditions (**Table 1** [18-22]). Each cell displays the average log2 fold change across donor replicates within the respective experiment. Color intensity reflects magnitude and direction of regulation (blue = upregulated; red = downregulated). Green stars denote genes that demonstrate consistent upregulation across multiple datasets, while red stars indicate genes showing reproducible downregulation. These annotations highlight directionally conserved remodeling patterns within ubiquitin-modifying enzymes. (βG, β-glucan; BCG, Bacillus Calmette–Guérin; PBMCs, peripheral blood mononuclear cells; LPS, lipopolysaccharide; YFV, yellow fever virus)

Together, these opposing expression patterns suggest selective remodeling of the ubiquitin network rather than uniform activation or repression, consistent with a coordinated shift in post-translational regulatory capacity.

### Comparison with RPMI-restimulated controls reveals selective persistence of ubiquitin signatures

To determine whether these transcriptional changes reflected trained immunity–specific reprogramming rather than generic inflammatory activation, trained samples were compared to RPMI-restimulated controls available in multiple datasets [20] - [22]. Normalization to these controls substantially reduced many baseline-relative expression differences (**Figure 3**), indicating that a portion of the observed signal arises from acute immune activation triggered by restimulation.

**Figure 3.**
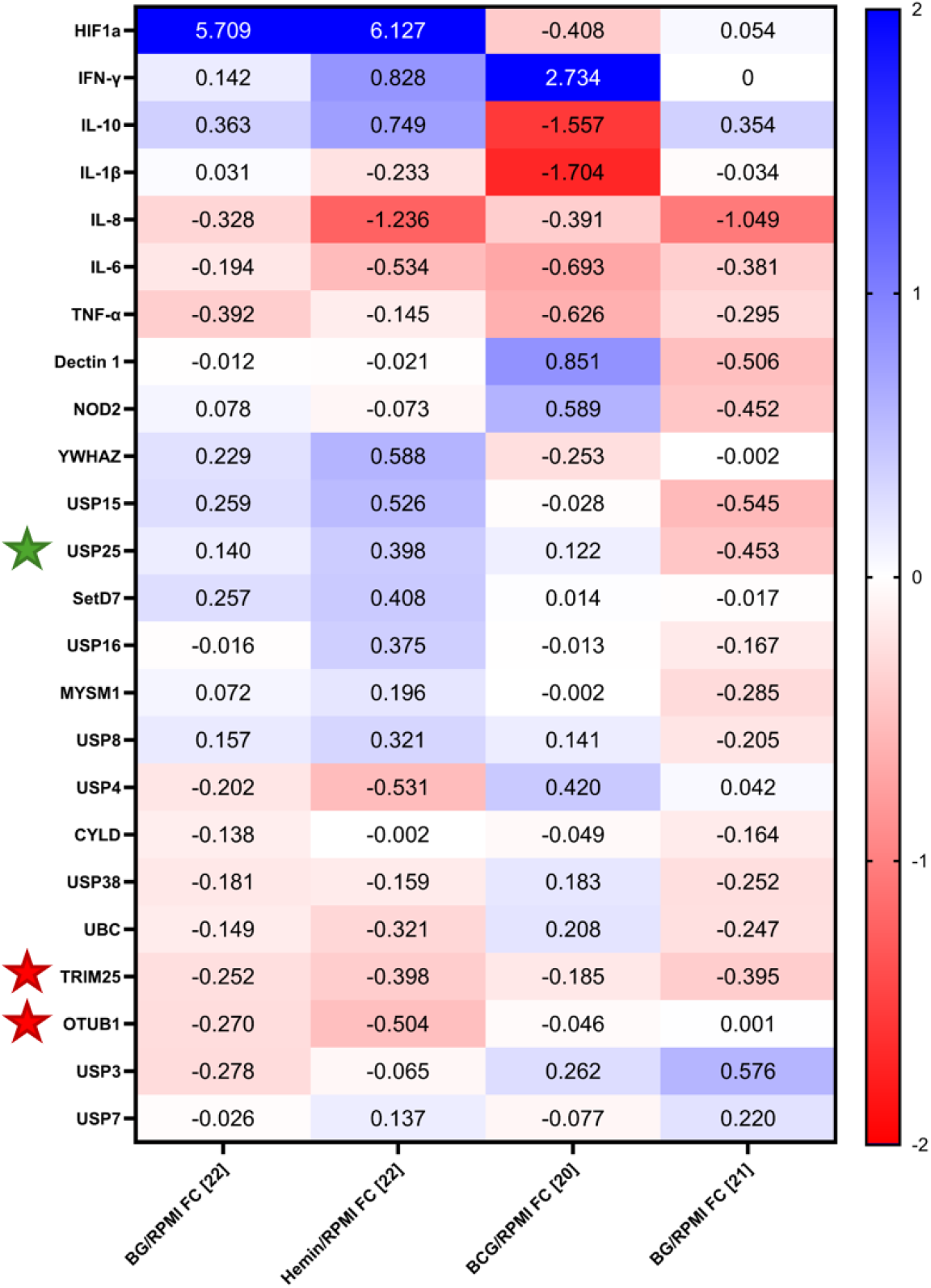
Heatmap of average log2 fold change in gene expression relative to RPMI- restimulated controls across trained immunity datasets. Rows represent individual genes and columns correspond to trained immunity conditions normalized to their respective RPMI restimulation controls (**Table 1** [20–22]). Each cell displays the average log2 fold change across donor replicates within the indicated dataset. Color intensity reflects magnitude and direction of regulation (blue = upregulated; red = downregulated). Green stars denote genes that retain reproducible upregulation following RPMI normalization, while red stars indicate genes that demonstrate consistent downregulation across conditions. These annotations highlight ubiquitin-modifying enzymes that maintain directionally conserved transcriptional signatures beyond acute restimulation effects.

However, USP25 remained consistently elevated relative to RPMI controls, distinguishing it from the majority of other genes analyzed. TRIM25 and OTUB1, while upregulated relative to baseline, displayed diminished or reversed changes when normalized to control conditions. The persistence of USP25 expression suggests that it may represent a stable transcriptional feature of trained immunity rather than a transient activation marker.

These findings support a model in which trained immunity induces selective stabilization of specific ubiquitin regulators while broader transcriptional responses remain largely stimulus dependent.

### Pathway enrichment analysis reveals coordinated engagement of ubiquitin regulatory networks

To evaluate whether individual gene changes converged on higher-order biological processes, Gene Ontology (GO) and KEGG pathway enrichment analyses were performed across ranked gene expression datasets. Multiple ubiquitin-associated functional categories were significantly enriched across independent training conditions, including pathways related to ubiquitin-like protein ligase binding (**Figure 4**).

**Figure 4.**
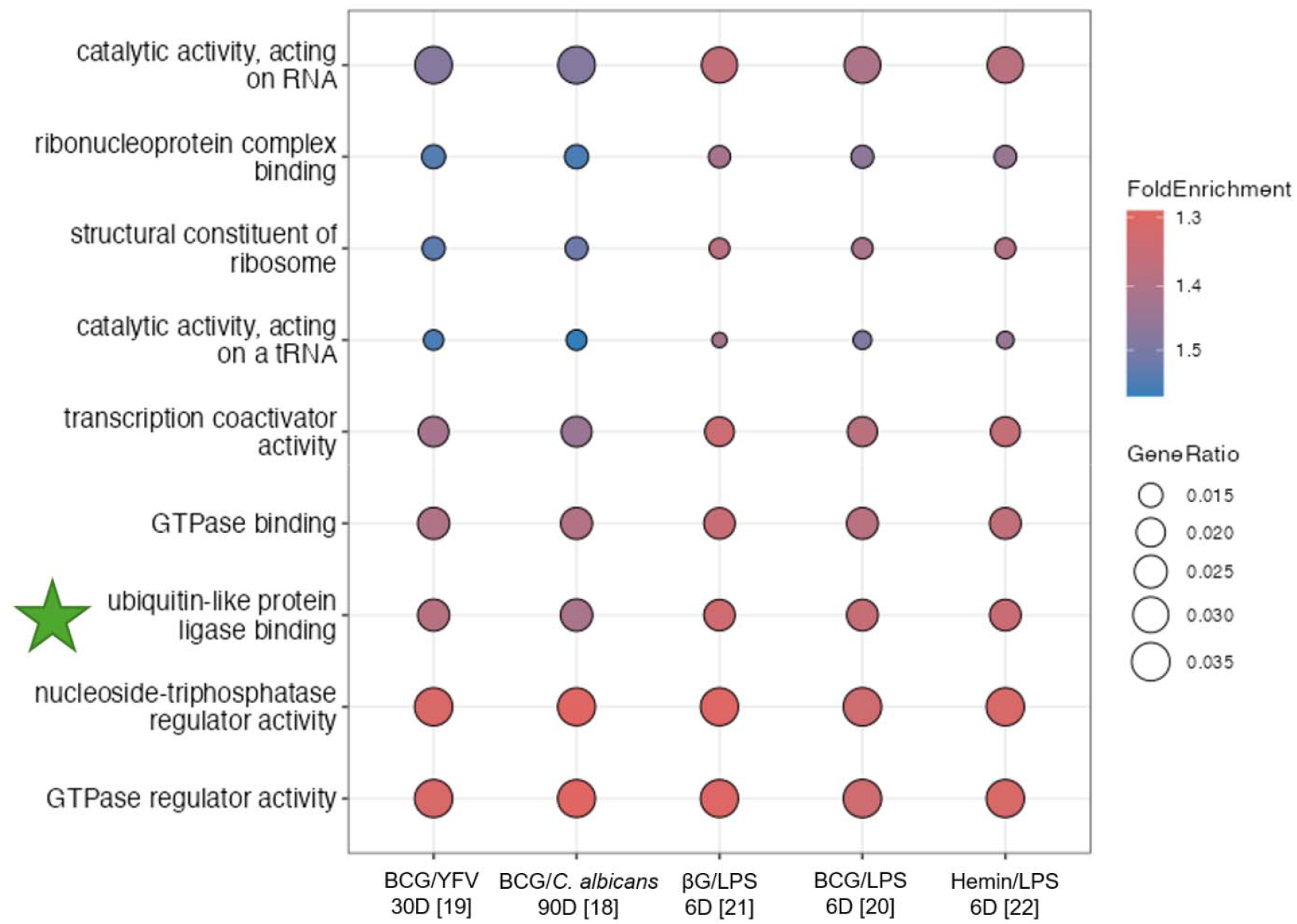
Pathway enrichment analysis across trained immunity datasets highlights reproducible engagement of ubiquitin-associated regulatory processes. Gene Ontology (GO) and KEGG enrichment analyses were performed on ranked gene expression profiles from independent trained immunity datasets. Each dot represents an enriched functional category; dot size indicates gene ratio, and color denotes adjusted p- value. Ubiquitin-related categories, including ubiquitin-like protein ligase binding, appear consistently across training conditions as indicated by the green star. Dataset identifiers and experimental annotations are provided in **Table 1**.

The reproducible enrichment of ubiquitin-linked processes across datasets indicates that transcriptional remodeling extends beyond isolated gene alterations and reflects coordinated reorganization of ubiquitin regulatory networks during trained immunity activation. These pathway-level findings independently reinforce the gene-specific observations (**Figure 2**) and support the hypothesis that ubiquitin signaling constitutes a systems-level component of innate immune memory.

### Functional integration of ubiquitin remodeling with immune signaling

Collectively, the transcriptional patterns and enrichment results suggest a dual regulatory strategy within the trained ubiquitin landscape. Upregulated enzymes such as USP25, OTUB1, and TRIM25 are primarily associated with antiviral readiness and controlled inflammatory signaling, whereas downregulated DUBs are linked to chromatin repression and attenuation of NF-κB activity.

This coordinated restructuring may promote a poised activation state characteristic of trained innate cells, enhancing responsiveness while preserving regulatory balance. The convergence of gene-level and pathway-level evidence supports the interpretation that ubiquitin signaling functions as an integrated modulatory axis within trained immunity.

## Discussion

### Trained immunity and ubiquitin remodeling

Trained immunity is an important aspect of innate immune response in myeloid cells that mediates an increased response upon pathogen re-exposure and serves as a form of non-specific memory [81], [83], [84]. While its regulation through metabolic and epigenetic remodeling has been extensively characterized, the contribution of post-translational modification systems such as ubiquitination has remained comparatively underexplored. In this study, we evaluated transcriptional patterns of deubiquitinating enzymes (DUBs) and E3 ubiquitin ligases across multiple trained immunity datasets to determine whether ubiquitin signaling exhibits coordinated remodeling during this process.

Our gene-level analysis identified consistent upregulation of USP25, OTUB1, and TRIM25 alongside downregulation of several chromatin- and cytokine-regulatory DUBs (**Figure 1 & Figure 2**). Importantly, pathway enrichment analysis independently revealed reproducible enrichment of ubiquitin-associated regulatory categories across datasets (**Figure 4**), indicating that these observations extend beyond isolated gene fluctuations and reflect broader transcriptional engagement of ubiquitin regulatory networks. Together, these results support the hypothesis that ubiquitin signaling forms an integrated component of the trained immunity landscape rather than representing incidental transcriptional noise.

### Ubiquitin signaling as a regulatory layer in trained immunity

The ubiquitin–proteasome system (UPS) functions not only as a degradation pathway but also as a dynamic signaling architecture integrating NF-κB, IRF3/7, and RIG-I/MAVS pathways [5] - [10]. The balance between ubiquitin ligases and DUB activity determines the amplitude and duration of immune signaling. The coordinated transcriptional changes observed here suggest that trained immunity is accompanied by restructuring of this regulatory axis.

Upregulated enzymes such as USP25, OTUB1, and TRIM25 are associated with antiviral readiness and controlled inflammatory signaling, acting on intermediates including TRAF3, TRAF6, and RIG-I [23] - [30], [50], [56]. Their repeated appearance across datasets, together with enrichment of ubiquitin-related functional categories, suggests that trained immune activation is coupled with reinforcement of ubiquitin-mediated regulatory capacity. Conversely, the suppression of DUBs linked to chromatin repression and NF-κB attenuation may transiently reduce inhibitory constraints, permitting enhanced transcriptional responsiveness upon secondary challenge [31] - [44], [46], [52].

Rather than a uniform shift toward activation or repression, these patterns imply selective tuning of ubiquitin architecture. This selective remodeling aligns with the functional hallmark of trained immunity: heightened responsiveness balanced by mechanisms that prevent uncontrolled inflammation.

### Dual role of ubiquitin remodeling

The dual nature of ubiquitin signaling is reflected in the opposing expression trends observed. USP25, which remains elevated even after normalization to restimulated controls (Figure 3), may represent a stabilizing checkpoint that preserves signaling equilibrium. Its known capacity to restrain excessive NF-κB activation while supporting type I interferon pathways is consistent with a regulatory role that enables robust but controlled immune memory [23] - [25].

In parallel, downregulation of USP3, USP4, USP7, USP16, MYSM1, and USP38 may transiently release repression of inflammatory or chromatin-associated pathways [31] - [33], [36] - [38], [46], [52]. Suppression of histone-linked DUBs in particular may increase promoter accessibility and reinforce epigenetic states associated with trained immunity [34], [35], [39] - [44]. These transcriptional shifts, viewed alongside pathway enrichment data, suggest that ubiquitin editing acts as a modulatory rheostat that shapes immune amplitude rather than serving as a binary switch.

### Crosstalk with metabolic and epigenetic remodeling

Trained immunity is defined by durable chromatin modifications and metabolic rewiring toward aerobic glycolysis [3] - [5]. Ubiquitin signaling intersects with both domains. Metabolite availability influences chromatin-modifying enzymes, while redox state and metabolic flux can alter DUB catalytic activity. The coordinated enrichment of ubiquitin-related processes suggests that ubiquitin remodeling is not an isolated phenomenon but may represent a parallel regulatory layer operating alongside metabolic and epigenetic adaptation.

Upregulation of antiviral-associated DUBs may stabilize signaling complexes during metabolic shifts, whereas suppression of chromatin-associated enzymes may facilitate enhancer accessibility. These relationships support a model in which ubiquitin architecture contributes to maintaining the transcriptionally poised state characteristic of trained innate cells.

### Ubiquitin signaling as a modulatory axis

Collectively, these findings support a framework in which ubiquitin signaling functions as a modulatory axis within trained immunity. Primary stimulation induces metabolic and epigenetic reprogramming accompanied by selective remodeling of ubiquitin networks. During the resting phase, persistent regulators such as USP25 may maintain equilibrium, preserving readiness without chronic activation. Upon secondary challenge, this reorganized ubiquitin architecture cooperates with open chromatin and altered metabolism to produce amplified but controlled immune responses.

In this context, ubiquitin-modifying enzymes operate as molecular rheostats that adjust signal intensity and duration. Integrating ubiquitin biology into the established metabolic– epigenetic paradigm expands the conceptual framework of trained immunity to include a complementary layer of post-translational regulation.

## Conclusion

This integrative analysis identifies coordinated transcriptional remodeling of ubiquitin regulatory networks as a recurring feature of trained immunity. The convergence of gene-level and pathway-level evidence suggests that ubiquitin signaling constitutes an additional regulatory layer within innate immune memory. By linking post-translational regulation to metabolic and epigenetic remodeling, these findings extend the current conceptual model of trained immunity and provide a framework for future mechanistic exploration.

A limitation to this study is that it relies on transcriptomic data sets that vary in donor characteristics, time points, and stimulation conditions, introducing heterogeneity. Moreover, transcript levels do not necessarily equate to enzymatic activity, and post-transcriptional regulation or ubiquitin turnover kinetics may alter functional outcomes. While enrichment analysis strengthens system-level interpretation, the results remain correlative. Future work should include functional perturbation and proteomic studies to determine whether the observed transcriptional patterns directly mediate trained immunity phenotypes.

## Methods

### Gene expression dataset selection

Publicly available transcriptomic datasets examining trained immunity were identified through the NCBI Gene Expression Omnibus (GEO) database. Inclusion criteria required: (1) primary exposure to a validated trained immunity stimulus (e.g., BCG, β-glucan, or hemin–β-glucan), (2) a defined resting interval, and (3) secondary stimulation with an immune challenge.

Datasets meeting these criteria and containing donor-matched baseline controls were selected for comparative analysis (**Table 1**). Only experiments performed in primary human monocytes, macrophages, or peripheral blood mononuclear cells (PBMCs) were included to minimize cross-species and cell line variability.

### Gene panel definition

A curated panel of ubiquitin-modifying enzymes was assembled prior to analysis based on literature-established roles in innate immune signaling, antiviral responses, NF-κB regulation, or chromatin remodeling (**Supplementary Table 1**). This targeted panel included deubiquitinating enzymes (DUBs) and E3 ubiquitin ligases with experimentally validated immune functions. Canonical trained immunity markers and inflammatory cytokines were analyzed in parallel as internal biological comparators (**Supplementary Tables 2–3**).

This targeted approach was used to evaluate hypothesis-driven transcriptional remodeling rather than perform an unbiased genome-wide screen.

### Differential expression analysis

Expression data were analyzed using the GEO2R interface to extract normalized gene expression values. Fold change was calculated by comparing trained/restimulated samples to matched baseline controls within each dataset. Log2-transformed fold change values were used for visualization and cross-dataset comparison.

Housekeeping genes YWHAZ and UBC were used as internal expression references to assess relative stability across conditions. Parametric paired t-tests were performed to evaluate statistical deviation from housekeeping gene expression (p < 0.05).

Because datasets originated from independent studies with differing preprocessing pipelines, analysis focused on within-dataset relative comparisons rather than cross-platform absolute expression values.

### RPMI control normalization

Where available, trained samples were additionally normalized to RPMI-restimulated controls to distinguish persistent training-associated transcriptional signatures from acute activation effects. Fold change was recalculated relative to RPMI controls using the same log2 transformation framework described above.

### Pathway enrichment analysis

To evaluate higher-order biological convergence, ranked gene lists from each dataset were subjected to Gene Ontology (GO) and KEGG pathway enrichment analysis using the clusterProfiler package in R. Enrichment was assessed using adjusted p-values to account for multiple hypothesis testing. Functional categories were considered significant when meeting standard false discovery rate (FDR) thresholds.

Ubiquitin-associated terms were interpreted in the context of reproducibility across independent datasets rather than isolated significance within a single experiment.

### Statistical considerations

Analyses emphasized reproducible directional trends across independent datasets rather than reliance on a single cohort. Because experimental designs varied across studies, the goal was to identify convergent transcriptional signatures rather than compute pooled meta-analytic effect sizes.

