## Supplementary Material for "Transcriptional remodeling of ubiquitin regulatory networks during trained immunity"

1. **Supplementary Tables**

**Supplementary Table 1.** A brief description of target ubiquitin-affecting molecules as well as their roles within the context of immune signaling.

| **Protein Name** | **DUB or E3 Ligase** | **Primary Targets** | **Immune Role** | **Reference** |
| --- | --- | --- | --- | --- |
| USP25 | DUB | TRAF3  TRAF6 | Type 1 interferon (IFN) response.  Reduce proinflammatory cytokine production. | [23] - [25] |
| USP16 | DUB | IKKβ  Histone H2A | NF-κB pathway induction  T-cell activation. | [39], [40], [43], [44] |
| USP8 | DUB | K63-linked ubiquitin chains on endosomes | Repress NF-κβ and Nrf2 signaling to prevent immune activation. | [45] |
| USP7 | DUB | MyD88  Raf-1 | Stabilizes a critical adaptor protein for TLR signaling and NF-κβ activation.  Regulates ERK1/2 signaling pathway. | [37], [46] |
| CYLD | DUB | MyD88  TRAF6  STING | Negative NF-κβ regulator in immune cells.  Important checkpoint in STING-mediated DNA sensing and antiviral response. | [47] - [49] |
| OTUB1 | DUB | UBC13  RIG-I | Promotes NF- κβ signaling via stabilizing UBC13.  Activates RIG-I-dependent antiviral signaling. | [29], [50], [51] |
| USP3 | DUB | RIG-I  MyD88 | Negative regulator for type I IFN signaling and NF-κβ activation.  Attenuates proinflammatory and antiviral response. | [31], [52] |
| MYSM1 | DUB | STING  RIP2  TRAF3  TRAF6 | Regulates antiviral and proinflammatory immune response through interaction with TRAFs, STING, and RIP2. | [34], [35] |
| USP15 | DUB | RIG-I | Promotes cytosolic DNA sensing and type I IFN amplification through cGAS-STING activation. | [53], [54] |
| USP38 | DUB | TBK1  KDM5B | Negatively regulates type I IFN signaling by editing active TBK1.  DUB activity coupled with the demethylase KDM5B to regulate H3K4 trimethylation and proinflammatory activity. | [41], [42] |
| USP4 | DUB | RIG-I  TRAF6 | Positive regulator of RIG-I activity and enhances NF-κβ activation by targeting TRAF6. | [32], [33], [55] |
| TRIM25 | E3 Ligase | TRAF2  TRAF6  RIG-I | Ubiquitinates RIG-I to enhance RNA virus signaling, IFN signaling, and modulate TRAF2/6 signaling. | [26] - [28], [56] |

**Supplementary Table 2.** A brief description of immune-related cytokines analyzed in this study.

| **Protein Name** | **Primary Function** | **Immunity Target** | **Reference** |
| --- | --- | --- | --- |
| IFN-γ | Involved in macrophage priming and activation.  Enhances antigen presentation. | Viral and bacteria | [57] - [59] |
| IL-10 | Suppress production of pro-inflammatory cytokines and inhibits antigen presentation. | Broad spectrum | [60] - [62] |
| IL-8 | Potent chemoattractant for neutrophils to infection and injury sites. | Bacterial | [63] - [65] |
| TNF-α | Promotes inflammation, induces apoptosis, and regulates other immune responses | Viral and bacterial | [66] - [68] |
| IL-6 | Induces acute phase proteins and mediates both pro- and anti-inflammatory responses. | Broad spectrum | [69] - [71] |
| IL-1β | Key mediator for inflammation and important lymphocyte activator. | Broad spectrum | [72], [73] |

**Supplementary Table 3.** A brief description of trained immunity markers analyzed in this study.

| **Protein Name** | **Primary Function** | **Trained Immunity Marker** | **Reference** |
| --- | --- | --- | --- |
| SetD7 | Methylates histones and transcription factors.  Epigenetic regulator for pro- and anti-inflammatory gene expression | Epigenetic marker | [74], [75] |
| NOD2 | Pattern recognition receptor (PRR) for bacterial markers.  Supports cytokine production via epigenetic changes in innate immune cells. | Epigenetic marker | [76] - [78] |
| HIF1α | Important regulator for hypoxia in immune cells.  Drives glycolytic shift in trained monocytes and macrophages via the mTOR-HIF1α pathway | Metabolic marker | [79] - [81] |
| Dectin-1 | Main β-glucan receptor.  Key initiator and coordinator for both innate and adaptive immunity. | Both metabolic and epigenetic marker | [82] |
